# Polygenic basis of strong and rapid flowering time response to environment perturbations in wild *Arabidopsis thaliana* population

**DOI:** 10.1101/2023.07.02.547444

**Authors:** Yan Ji, Yu Han, Yifei Dai, Fan Hao, Xiao Feng, Qipian Chen, Ran Hao, Zhiqiang Chen, Wei Zhao, Wenjia Zhang, Huan Si, Yanjun Zan

**Affiliations:** Tobacco Research Institute, Chinese Academy of Agricultural Sciences, Qingdao, CN-266000, China; Key Laboratory for Bio-Resource and Eco-Environment of Ministry of Education & Sichuan Zoige Alpine Wetland Ecosystem National Observation and Research Station, College of Life Science, Sichuan University, Chengdu, CN-610065,China; Biostatistics Department, School of Public Health, University of Michigan, AnnArbor, Michigan, 48105, USA; College of Forestry, Northwest Agriculture and Forestry University, Yangling, CN-712100, China; Greater Bay Area Institute of Precision Medicine (Guangzhou), Fudan University, Guangzhou, CN-511400, China; Shenzhen Branch, Guangdong Laboratory of Lingnan Modern Agriculture, Genome Analysis Laboratory of the Ministry of Agriculture and Rural Affairs, Agricultural Genomics Institute at Shenzhen, Chinese Academy of Agricultural Sciences, Shenzhen, 518120, China; Umea□ Plant Science Centre, Department Forest Genetics and Plant Physiology, Swedish University of Agricultural Sciences, SE-90183 Umea□, Sweden; Department of Ecology and Environment Science, Umeå Plant Science Center, Umeå University, Umeå, SE-90187, Sweden

**Keywords:** Flowering time, Plasticity, climate change, Polygenic, Genotype by environment interactions

## Abstract

Despite the importance in understanding the impact of climate change, the genetics of rapid response to changing environments and its role in adaptive evolution remains elusive. Here, we studied flowering time response to environment changes using 514 *Arabidopsis thaliana* worldwide accessions with re-sequencing genomes and flowering time measurements from ten unique environments with variable temperature, drought, daylight and competition stresses. We revealed a polygenic basis of flowering time mean and plasticity, underpinned by 52 mean and plasticity QTL. Widespread interaction between mean QTL, polygenic background and surrounding environments considerably altered the amount of additive genetic variance and allelic effects of detected QTL. This caused variability in phenotype plasticity and across environment variation in genetic variance, resulting in rapid flowering time response to environment perturbations. In addition, the plastic alleles showed a higher correlation with the environment factors than that from randomly sampled alleles, suggesting a potential role in climate adaptation. We therefore proposed a polygenic interaction model, whereby large effect QTL and polygenic background simultaneously interacted with the surrounding environment, underlying rapid response to changing environments. Results from our study thus provided deeper insights into the genetics of plasticity, with potential benefit in genomic selection of crops in heterogeneous environments and predicting changes in species distribution and the evolutionary trajectory of wild populations.

## Introduction

Organisms display the ability to rapidly respond to changes in surrounding environment without requiring changes in their genetic makeup, namely phenotype plasticity or genotype by environment (G×E) interaction (Price *et al*., 2003). Nature environments vary spatially and temporally due to shifts in biotic or abiotic factors. Such variability in surrounding environments always creates a mismatch between genotype and environment within a generation. This is a timescale that not even rapid evolution could take place. Plasticity allows organisms rapidly change their phenotypes to align phenotype with the surrounding environments, therefore, could have a profound impact on evolution and an increased understanding of the genetic basis of plasticity is highly relevant in better our understanding of the maintenance of genetic variation (Gillespie & Turelli, 1989), complex trait variation (Forsberg *et al*., 2017), genomic selection of crops in heterogeneous environments (Jin *et al*., 2023) and predicting changes in species distribution and the evolutionary trajectory of wild populations (Kelly, 2019).

Decades of experimental evolution and artificial selection experiments have revealed remarkable selection responses from shifts in frequencies of many minor effect alleles, aka standing genetic variations, concluding a polygenic basis for long-term response to selection (Sheng *et al*., 2015). For decades, efforts have been made to study plasticity (Finlay & Wilkinson, 1963; Gollob, 1968; Jiang & Zeng, 1995; Malosetti *et al*., 2013) and dissect the underlying QTL (El-Soda *et al*., 2014; Rauw & Gomez-Raya, 2015; Kusmec *et al*., 2017, 2018; Li *et al*., 2019; Schneider *et al*., 2020). Evidence suggests that phenotype plasticity in *Arabidopsis thaliana* and maize likely (Gage *et al*., 2017; Jin *et al*., 2022) involves a large number of loci. However, a detailed understanding on specific genetic variants and genes that contribute to variation in plasticity is lacking. This limited our ability to understand a few key questions. First, what is the relationship between the genetic basis of phenotype mean and phenotype plasticity? And what is the role of mean quantitative traits locus (QTL) and plastic QTL in short- and long-term responses to environment changes? Second, what proportion of alleles could alter their effects across environments, and what is the relative role of QTL by environment interaction and interaction between polygenic background and surrounding environment in variation in across environment phenotype/genetic variance, aka cryptic genetic variation? Last but not least, how to fit plasticity in the evolutionary biology theory in the context of adaptive evolution.

To address these questions, we evaluated the genetic basis of flowering time mean and plasticity in response to environment changes using 514 *Arabidopsis thaliana* worldwide accessions with re-sequencing genomes and flowering time (FT) measurements from ten unique environments. We revealed that the genetic basis of flowering time mean was a dynamic property of the growth environments, whereby environment change considerably altered the amount of additive genetic variance and allelic effects of detected QTL. In addition, we demonstrated that aggregated changes from the allelic effects of large effect QTL and standing genetic variations made a remarkable impact on the phenotypic response to environment changes. The plastic alleles showed a higher correlation with the environment than that from randomly sampled alleles, suggesting a potential role in climate adaptation. Results from our study thus provided deeper insights into the genetics of plasticity, with potential benefit in genomic selection of crops in heterogeneous environments and predicting changes in species distribution, community composition and the evolutionary trajectory of wild populations.

## Results

### Study design

We reanalysed two public datasets with measured flowering time from 514 *Arabidopsis thaliana* worldwide accessions (Alonso-Blanco *et al*., 2016; Exposito-Alonso *et al*., 2019). The first dataset [13] involved two common gardens at Tu□bingen and Madrid, where drought and competition stress was simulated by manipulating water availability and plant density at each site, resulting in eight unique environments (The name of each environment is Illustrated and abbreviated in Fig 1A). The second dataset [14] includes two flowering time measurements, one at 10 C° and a second one at 16 C°, under greenhouse conditions (Fig 1A). All the flowering time measurements were taken as the mean of four to eight replicates, referred to as flowering time mean (FTm) measurements hereafter. Altogether, our study included FTm measured from ten unique growth conditions (Fig 1A), with corresponding abbreviations illustrated in Fig 1A.

**Fig 1.**
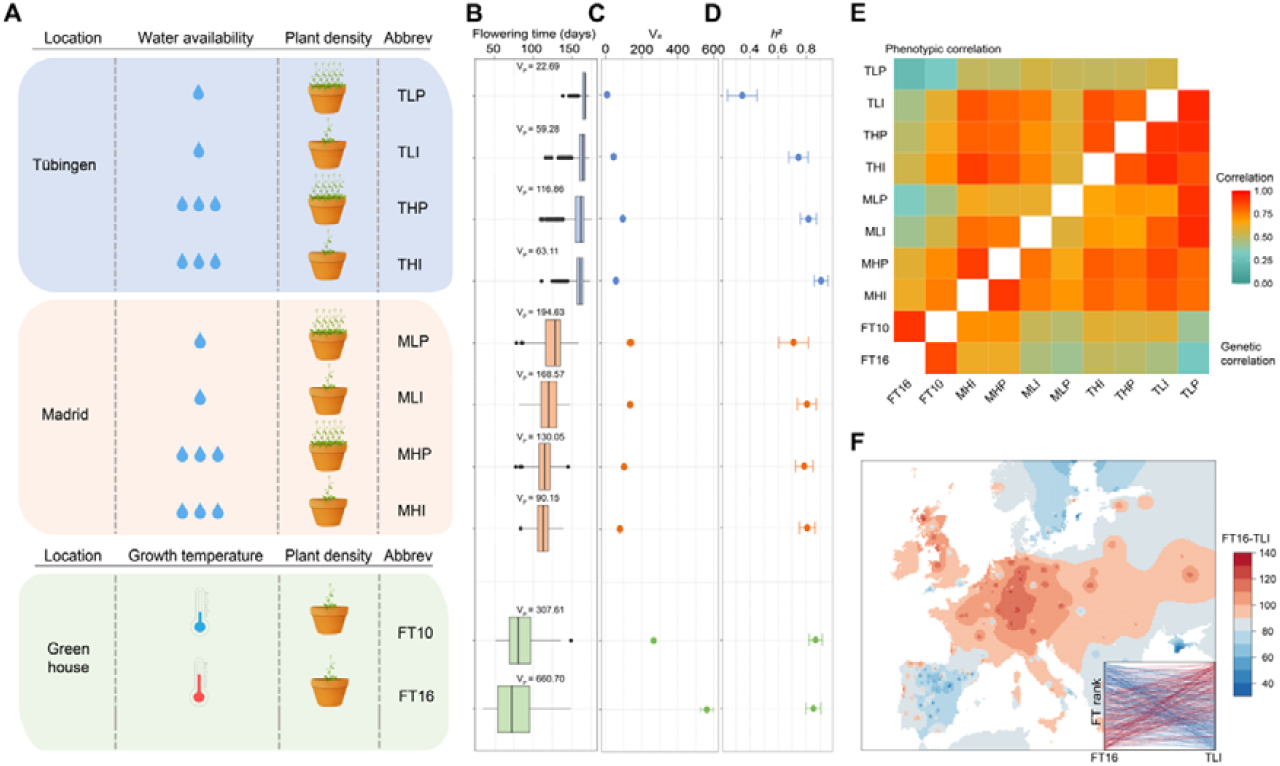
**A)** A schematic illustration of the ten studied environments. **B)** Boxplot of flower time mean measured at ten studied environments. **C)** Estimated additive variance at each environment. **D)** Estimated narrow-sense heritability (*h*^*2*^) for within replicates FT means at each environment. Each dot represents a point estimate of *h*^2^ indicated on the y-axis and the corresponding environment was labelled in the x-axis. Standard errors were plotted as error bars. **E)** Upper triangular: Pairwise Spearman rank correlation among FTm measured under ten environments. **E)** Lower triangular: Pairwise genetic correlation among FT measured under ten environments. **F)** Reaction norm of flowering time between FT16 and TLI. Each line represents an inbred line, and the lines connect the rank of flowering time from two environments.

### Extensive variation in Arabidopsis thaliana flowering time mean and plasticity

FTm displayed continuous variation (Fig 1B), being the shortest at 73.67 days (FT16) and the longest at 165.23 days (TLP). Both drought and competition-induced stress changed the mean and phenotypic variance of FTm (V_p_; Fig 1B) across environments. Compared to environments involving single stress, such as drought (XHI Vs XLI, X stands for either Tu□bingen or Madrid, H and L stand for high or low water availability), competition (XHI Vs XHP, or XLI Vs XLP, I and P stand for being planted as individual or a population of 30 individuals in a pot), or growth climate (TYI Vs MYI, or TYP Vs MYP, Y stands for either H or L), combinations of two or multiple stresses yielded a larger difference in both mean and variance of FTm, resulting in decreased pairwise correlations (Fig 1E), even though the FTm from each environment were highly heritable (*h*^*2*^ ranged from 0.34 to 0.90, with a median equal to 0.80; Fig 1D).

There was a widespread reaction norm for FTm. For example, when the environment changed from TLI to FT16, accessions from the central of Europe increased flowering time considerably while others from a large part of Sweden and Spain either remained stable or decreased rank drastically, leading to variation in flowering time plasticity (FTp, Fig 1E). We, therefore, quantified FTp using two types of metrics. The first type involved three measures including Principle component (PC), Finlay–Wilkinson Regression (FWR), and across environment variance in rank transformed FTm (VarR) quantifying the overall flowering time plasticity (OP) across all environments. The second metric was the pairwise FTm differences for all possible pairs of environments, quantifying specific changes across paired environments, resulting in a total of 45 specific plasticity (SP) measures. Almost all the plasticity measurements were continually distributed with intermediate to high narrow sense heritability (0.10-0.81, median = 0.61, Figure S1). In addition, the 48 plasticity measures, displayed variable correlation from 0.02 to 0.88 (Figure S2), indicating novel environments induced plastic response were often environment specific. This will be explored further in the following sections.

### Environment perturbation buffer and release additive genetic variation

There was moderate variation (0.71-0.90; TLP was excluded as hundreds of plants were stressed to death; Fig 1D) in the estimated narrow-sense heritability from one environment to another. Consistent with this, the amount of additive genetic variation (V_a_) estimated as V_p_ times *h*^*2*^ changed considerably from 44.00 (TLI) to 561.46 (FT16) (Fig 1C). Such changes in V_a_ was likely caused by either altered genetic effects from one or multiple QTL or turning on and off of a set of QTL, indicating that the genetic architecture of FTm was dynamic across the ten environments. This was supported by the variation of pairwise phenotypic/genetic correlation estimated from a multivariate mixed model analysis treating FTm from individual environments as independent traits (Fig 1E; Materials and methods). The estimated genetic correlation between environments was as low as 0.56 (outlier TLP was excluded), indicating considerable changes in breeding values from many individuals. Altogether, these results demonstrated that there was a considerable change in the genetic regulation of FTm across environments, which altered the amount of genetic variance and *h*^2^ for FTm in each environment. In the following sections, we will dissect the genetic basis underlying variation of FTm and FTp, and trace the origin of genetic variation that arose from environment change.

### Polygenic architecture underlying Arabidopsis thaliana flowering time mean and plasticity variation

A total of 52 QTL were identified for FTm and FTp using genome-wide association analysis (GWA), including ten flowering time mean (FTm), four overall plasticity measures (OP) and 45 specific plasticity measures (SP). This included 37 QTL for FTm, 18 QTL for SP and 15 QTL for OP with variable degrees of overlaps, indicating a shared genetic basis between FTm and FTp (Fig 3A/C, Table S1, Manhattan-/QQ-plots are available in Figure S3, Materials and Methods). By comparing the genetic effects of a QTL on chromosome 1:24 339 228 bp detected for a subset of the environments, we found that the genetic effect of this QTL changed from -2.72 ± 0.42 days (p = 9.04 × 10^−11^) for TLI to -3.67±1.22 days (p =2.62×10^−3^ Fig 3 A-C/E) for FT16, and this QTL is detected with association with OP (PC1, p =1.23×10^− 9^), suggesting that changes in the magnitude of genetic effects on flowering time mean may have contributed to variation in flowering time plasticity.

Since the number of QTL detected for each FTm/FTp measure at genome-wide significance threshold varied from 1-20 (Fig 2B), we tested whether the effect of additional QTL, detected for FTm at other environments but not focal environments, could explain significant genetic variance as a group. Results showed that although not every QTL was individually associated with each FTm, together they were making a significant contribution to all the FTm or FTp traits, suggesting a polygenic basis underlying flowering time mean and plasticity (P-values are available in Table S2; Materials and methods). Hereby, the genetic architecture of the FTm and FTp traits was defined to include all 52 QTL that together explained from 10.16% to 72.04% (median= 45.13%) of the within-environment phenotypic variance (Table S3).

**Fig 2.**
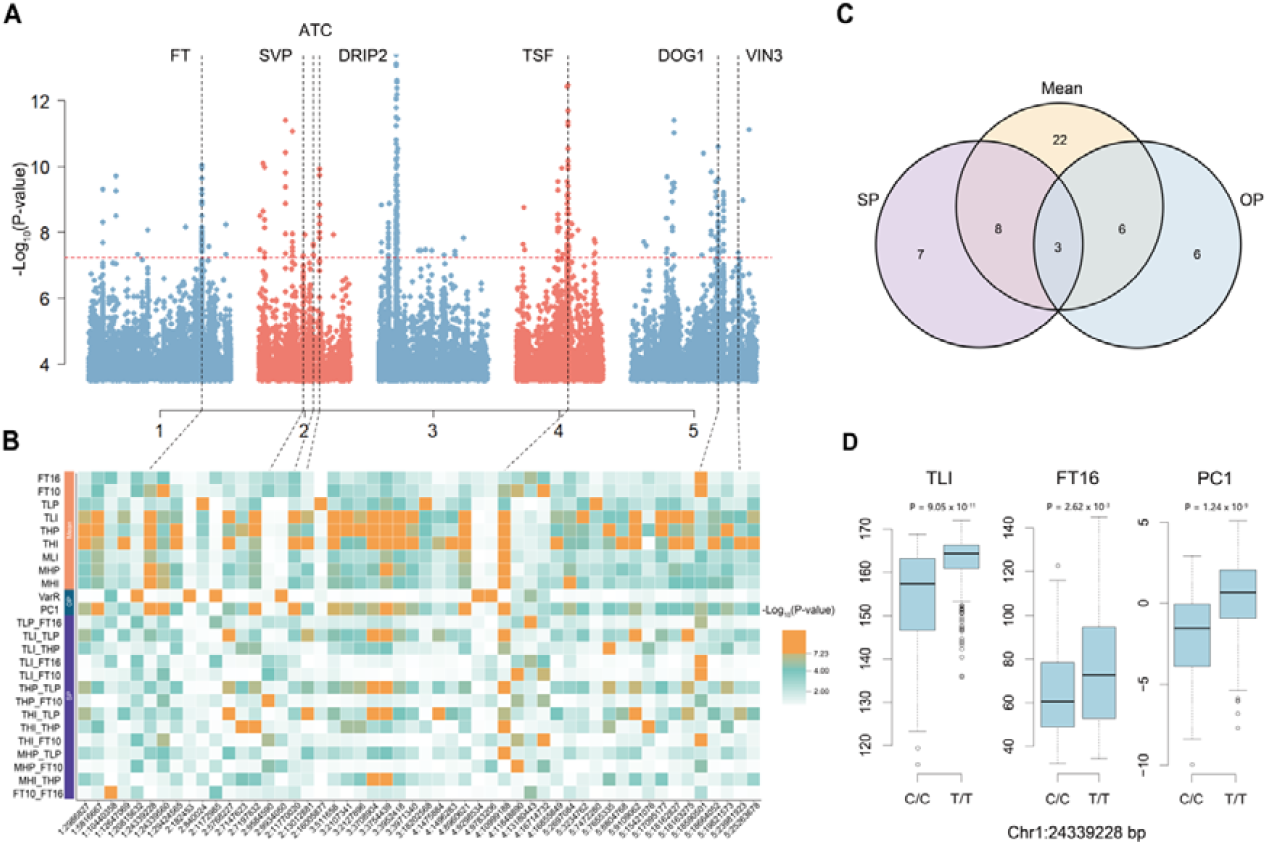
Summary of the GWA results for flowering time mean and plasticity. **A)**. Joint Manhattan plots overlaying results from 58 GWA scans, where 10 FTm and 48 FTp measurements were analysed, and -log10 (p values) were overlaid on panel A. The red horizontal dashed line indicates the Bonferroni corrected genome-wide significant threshold, and QTL peaks surrounded by known candidate genes were marked using dashed vertical grey lines. **B)** A heatmap illustrating the p values of 16 SNPs detected for more than two traits. Each cell represents the -log10 (p-value) of a particular SNP (x-axis) associated with a specific trait (y-axis on the right). **C)** Venn diagram illustrating the overlap of QTL detected for the 3 types of flowering time measurements. **D)** Genotype-to-phenotype maps for FT16, TLI and OP at chromosome 1: 24 339 228, ’*’ means p-value < 0.05 but greater than genome wide significant threshold and ‘**’ means p value smaller than the genome wide significant threshold.

**Fig 3.**
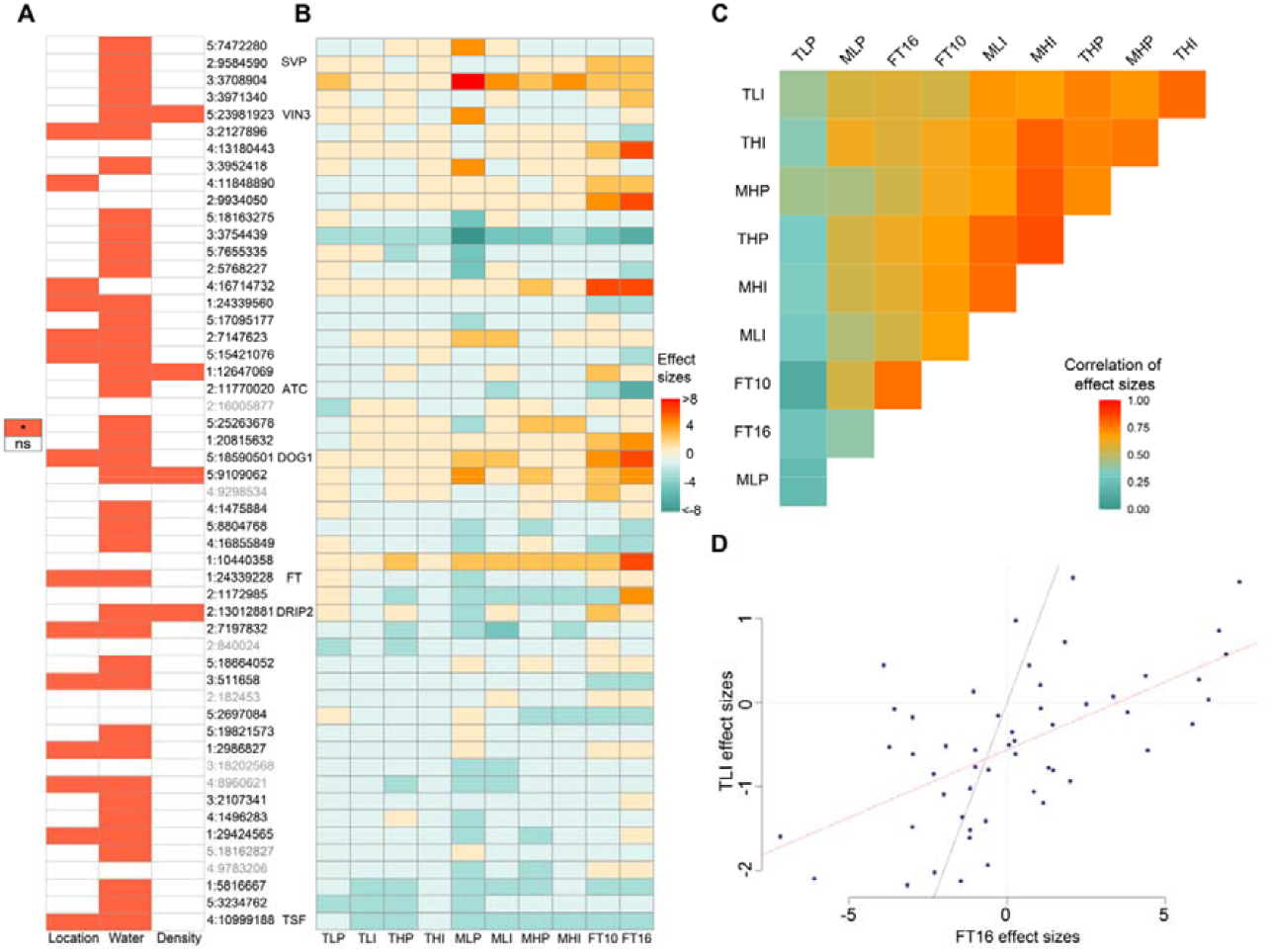
Variation of genetic effects across-environment. **A**) A heatmap illustrating whether a QTL is interacting with location, water availability or planting density. **B**) A heatmap illustrating the additive genetic effects (y-axis) from 52 QTL on FTm from ten environments (x-axis). **C)** Pairwise correlation in the 52 additive effects from ten environments. **D)** Correlation between the additive genetic effects for the 52 loci estimated for accessions grown at two extreme environments, FT16 and TLI. The red line indicates the regression line from linear regression.

As reported in previous studies (Atwell *et al*., 2010), linkage disequilibrium was too extensive to directly pinpoint the causative genes or variants in the 1001 Genome dataset. Nevertheless, we found a few flowering time regulators *FT, SVP, ATC, DRIP2, TSF, FLC, DOG1, and VIN3*, near the vicinity of the top associated SNPs (Schwartz *et al*., 2009; Wang *et al*., 2012; Hiraoka *et al*., 2013; Méndez-Vigo *et al*., 2013; Li *et al*., 2014; Huo *et al*., 2016) (Table S1). Some of these genes, such *FT, TSF* and *DOG1*, were detected in most of the environments while the remaining candidates were environment specific.

### A large proportion of the detected QTL interacted with environment

38 QTL (75%) were only detected for one of the mean or plasticity measurements, while 9 QTL were shared more than 5 times (Table S1). Such variation in QTL being detected in some measures but not others could result from either difference in detection power (significance threshold being too conservative) or turning on/off of genetic effects across the environment due to genotype-by-environment interaction. We, therefore, first compared the overlap under a lenient significance threshold correcting for the 52 QTL. In this case, more than 60% of the detected QTL were unique to one or only a few environments, indicating a proportion of the QTL had environment-dependent effects. This was supported by further analysis testing genotype by environment interactions for the 52 QTL one at a time using a multivariate mixed model (Materials and Methods). In total, 41 (80.77%) QTL displayed significant genotype by interaction, with 14, 39 and 4 QTL interacting with growth site, drought and plant density, respectively (Fig 3A, Table S4). For QTL displaying significant genotype by environment interaction, their allelic effects were relatively stable at most of the environments with one or very few environments showing significantly increased/decreased allelic effects, suggesting that novel environment activated standing genetic variations that were neutral or nearly neutral under most of the environment condition. Between the two environments with the longest (TLI) and shortest (FT16) FTm, we observed a change of effect sizes across most of the QTL (Fig 5D). These results indicated that the difference in FTm across the environment was likely the result of altered allelic effects across many loci. This will be explored further in the following sections.

### Across environment variation in genetic variation (aka cryptic genetic variation) arise from interaction between multiple QTL, polygenic background and environment

As has been mentioned above, stresses introduced by drought, planting density, and overall growth climate, resulted in large differences in phenotypic and additive variance across the ten environments. When we treated FTp as a trait and measure it as the difference in FTm between two environments, highly heritable quantitative variations were observed for all the plasticity measurements and many QTL were detected for FTp, with a large proportion of the plasticity QTL showed environment-dependent genetic effects. In this section, we will evaluate the role of genotype-by-environment interaction in the variation of V_p_ and V_a_ across environment (aka cryptic genetic variation), and their contribution to the variation of FTm and FTp, focusing on the contribution from the detected QTL and polygenic background.

#### Contribution from the 52 QTL and polygenic background to variation of phenotypic and additive variance across the ten environments

Here, we explored whether across environments differences in phenotypic/genetic variance could be explained by the conditional effects from the 52 QTL, polygenic background, or both. FTm from each environment were partitioned to contribution from 52 QTL and polygenic background by fitting a mixed model with an intercept, fixed effects of allelic effects from the 52 QTL, and polygenic effects with identity-by-state (IBS) matrix as covariance structure. This analysis was repeated for the ten environments one at a time, where, for each plant, the aggregated effect from 52 QTL and polygenic background was estimated as the sum of the allelic effects from the 52 QTL and the breeding values, respectively. For each environment, variation in the aggregated effects from the 52 QTL (V_52QTL_) and polygenic effects (V_pg_) was then calculated as the within-environment variance of corresponding estimates across the 514 individuals. Figure 4 A/B illustrated a significant correlation between V_52_ and V_p_ as well as V_Pg_ and V_p_, indicating a joint contribution from 52 loci and the polygenic effects to the across-environment variation in V_p_ and V_a_. However, the contribution from 52 QTL was, on average, 2.11 times larger than that from the polygenic background (Fig 4F), indicating the effects from large effect QTL were more important.

**Fig 4.**
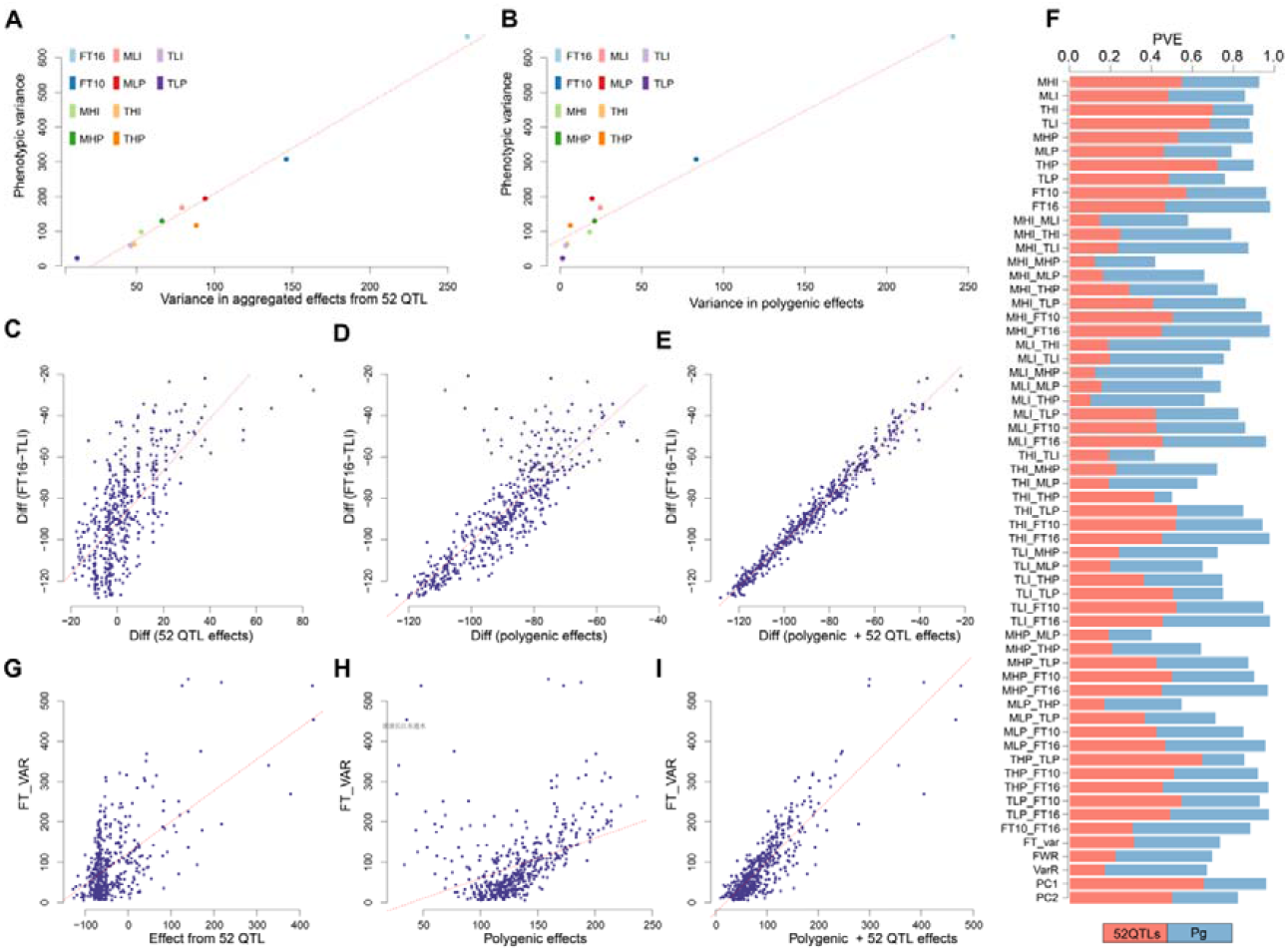
Genotype by interaction and origin of across environment phenotypic/genetic variation. **A)** Scatter plot for variance in aggregated effects from 52 QTL (x-axis; V_52_) and phenotype variance (y-axis; V_p_). Each dot represents an environment with the corresponding V_52_ on the x-axis and V_p_ on the y-axis. **B)** Scatter plot for variance in the polygenic background (x-axis; V_pg_) and phenotype variance (y-axis; V_p_). Each dot represents an environment with the corresponding V_pg_ on the x-axis and V_p_ on the y-axis. **C)** Relationship between aggregated effects from 52 QTL and the specific flowering time plasticity measured as the difference between FT16 and TLI. **D)** Relationship between aggregated effects from the polygenic background and the specific flowering time plasticity measured as the difference between FT16 and TLI. **E)** Relationship between aggregated effects from polygenic background plus the 52 QTL and the specific flowering time plasticity measured as difference between FT16 and TLI. **F)** Stacked bar plot illustrating the relative contribution from V_52QTL_ (tomato) and V_Pg_ (light blue) to across environment variance of flowering time. **G)** Relationship between aggregated effects from 52 QTL and the overall flowering time plasticity measured as within-line variance across 10 environments. **H)** Relationship between aggregated effects from the polygenic background and the overall flowering time plasticity measured as within-line variance across 10 environments. **I)** Relationship between aggregated effects from polygenic background plus the 52 QTL and the overall flowering time plasticity measured as within-line variance across 10 environments.

#### Contribution from the 52 QTL and polygenic background to variation in flowering time plasticity

Next, we explored the contribution of the 52 QTL and polygenic background to variation in SP and OP. First, we focused on SP measured as the pairwise difference in FTm, quantifying the flowering time plasticity changing from one particular environment to another. FTm from each environment was first partitioned to contribution from the 52 QTL (Eff_52_) and polygenic background (Eff_Pg_) one at a time as described above (Materials and Methods). Then, we evaluated how the difference in Eff_52_ (Abbreviated as Diff_52_, where 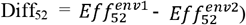 and Eff_pg_ (Abbreviated as Diff_pg_, where 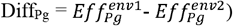) contributed to the difference in FTm (Abbreviated as Diff_FT_, where Diff_FT_ =FT^env1^ -FT^env2^; Materials and methods). We first illustrated results using FT16 and TLI, where FTm displayed the largest difference, and then extended the results to all the remaining pairs. As illustrated in Fig 1 B-C/E, there was extensive genotype by environment interaction between these two environments and the difference in FT16 and TLI showed continuous variation with narrow sense heritability equal to 0.81. Variation in Diff_FT_ between FT16 and TLI could be attributed to both Diff_52_ (Fig 4C) and Diff_pg_ (Fig 4D), where the contribution from 52 QTL was nearly equal to (0.94 times) that from the polygenic background in this particular case. Across all 45 pairwise combinations of FT plasticity measurements, 8.62% to 63.47% (median = 43.38%; Fig 4F) the variance in specific plasticity was explained by Diff_52_, which is 2.60% more than that form polygenic background (median = 40.78%; Fig 4F).

Second, we focused on OP measured as the variance of FTm across the ten environments. Following the same procedure described above, we partitioned the variance of OP measurements into contribution from the 52 QTL (Eff52) and polygenic background (Effpg). Most of the variance in OP was explained by the variation in polygenic effects (median =41.81%), which is 10.43% more that from the 52 QTL (median =31.38%). Altogether, these results suggested a polygenetic architecture of FTm and FTp with relatively larger contributions from polygenic effects that that from a few large effect QTL. Genotype by environment interaction between large effects allele and polygenic background together altered the genetic architecture, resulting in variation in phenotypic plasticity and across environment phenotypic and genetic variance.

### A model for strong and rapid response to environment perturbation and its implication in adaptive evolution

Based on the results presented above, we propose a model underlying the strong and rapid response to environment perturbation unifying genotype by environment interaction, the polygenic architecture of phenotype mean and plasticity (Fig 5). At individual environment, a subset of the QTL has detectable effects (highlighted as triangular, Fig 5A) at genome wide significance threshold, while others were buffered (highlighted as circles, Fig 5A) with genetic effects that were too small to be detected at stringent significance threshold. Meanwhile, the remaining loci across the genome were making minor contributions (aka polygenic effects) that were individually not detectable but together can explain up to 50 % of the phenotype variation. Upon environment change, a subset of the QTL will be deactivated or show considerably decreased effects. Meanwhile, the remaining loci across the genome would likely experience subtle changes in their effect sizes. However, the aggregated effects from large effect QTL and polygenic background for each individual could change considerably, generating remarkable phenotype change in response to environment perturbation. This type of interaction between a massive number of genes, aka polygenic architecture, and environment stimulators facilitate strong and rapid response to environment change without requiring any new mutation or shift in the frequencies of many alleles.

**Fig 5.**
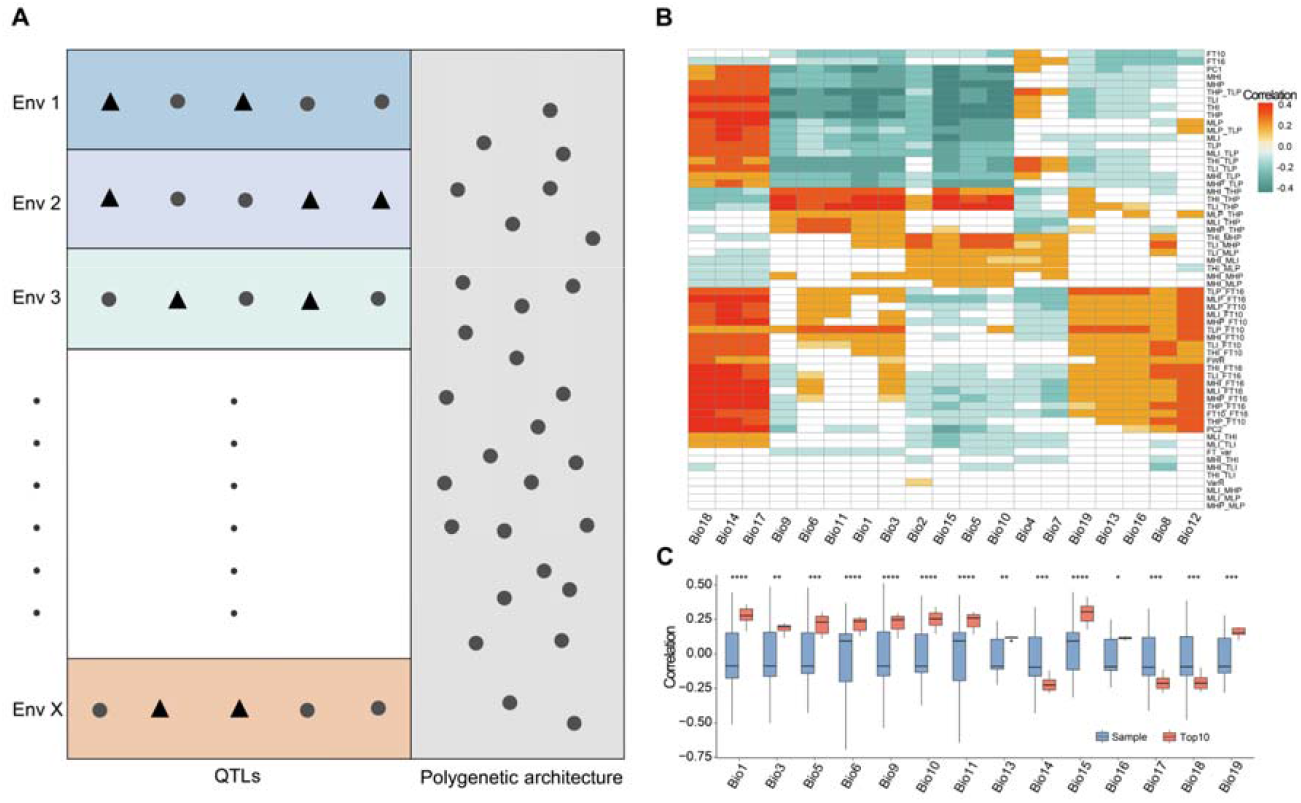
A unified framework for strong and rapid phenotype response to environment perturbation, and its role in climate adaptation. **A)** Schematic illustration of a model describing the genetic basis of for strong and rapid phenotype response to environment perturbation unifying polygenic architecture of phenotype mean and plasticity variation, genotype-by-environment interaction. **B)** A heatmap illustrating the correlation between FTm/FTp and environment variables obtained from the world climate database (http://www.worldclimate.org). Cells with pairwise correlation test p values above 0.05 were marked with white colour. **C)** Boxplot contrasting the Pearson correlations between the top ten percent plastic alleles (QTL with p values ranked at the top 10% in interaction test, tomato colour) and environment factors, and correlations between randomly sampled QTL (skyblue) and environment factors.

We hypothesise that such a dynamic genetic architecture with the ability to tune the genetic effects from large and small effect QTL in response to environment change likely played a role, at least to some extent, in adapting to a fluctuating environment. This was supported by the observation that SP and OP measurements were significantly correlated with environment variables at their naive range (Fig 5 B), and the top 10% plastic alleles (QTL with p values ranked at the top 10% in QTL×E test) showed higher correlation with environment variables than that form randomly sampled QTL (Fig C). Given that plasticity and plastic QTL are not independent of the trait mean and mean QTL, thereby, selection on plasticity and plastic QTL is likely constrained by their mean effects.

## Discussion

A central question in evolutionary quantitative genetics is the genetic basis of plasticity and whether it contribute to adaptive evolution. Here, we investigate these questions by quantifying the level of phenotype plasticity, dissecting the underlying genetic variants, studying their genetic effects across environments and their association with environment factors.

### Flowering time plasticity is property of the specific environment perturbation

Quantifying variation in flowering time plasticity under natural environments presents significant challenges due to complex, multivariate reaction norms from multiple environment factors. Our study overcome this by simulating drought and competition within each common garden, making it possible to study specific environment factor induced plasticity. The 48 specific plasticity measurements quantifying plasticity from one specific environments to another displayed variable correlation from 0.02 to 0.88 (Fig S2), indicating novel environments induced plastic response were often environment specific. Therefore, we adapted two matrix, specific plasticity and overall plasticity, to gain a better overview about the degree of plasticity. Although such multivariate reaction norms present a great challenging in comprehensively understanding the genetic basis of plasticity, they are likely making the populations more resilient to environment change, offering more opportunities to adapt to future challenges.

### Variation in plasticity is a result of the dynamic genetic architecture of FTm due to widespread interactions between environment

We demonstrated that the genetic basis of FTm variation is a dynamic property of the surrounding environments. Changes in environment cues activated, deactivated or altered the genetic effects of many large effects mean QTL and minor effects alleles form polygenic background. Their aggregated effects varied significantly across environments, resulted rapid response to environment perturbation. This detailed dissection of genetic basis of phenotype plasticity connects and unifies many previous coined terms in this field, such as reaction norm (Nicoglou, 2015), G×E (Nicoglou, 2015), plasticity (Bradshaw, 1965), canalization (Waddington, 1942), providing insights into the genetic mechanisms of rapid response to environment changes.

### Potential role of plasticity and plastic alleles in ecological adaptation

Currently, there are three non-mutually exclusive hypothesis on how plasticity could adaptation. First, plasticity could facilitate adaptation by promoting phenotypes close to optimal in novel environments, thereby buying time until adaptive evolution could occur (Merilä & Hendry, 2014) (the “buying time hypothesis”). Second, plasticity could facilitate adaptation by exposing cryptic genetic variation to selection (Levis & Pfennig, 2016) (the “Plasticity led evolution hypothesis”). Last, plasticity could form an alternative inheritance system on which adaptive evolution could act. Detailed dissection of the genetic basis of plasticity in our study supports that those hypotheses are not mutually exclusive, and were inter-connected. First of all, upon environment change, rapid changes in the allelic effects of multiple QTL did expose additional genetic variation to selection and thereby buying time for new mutation to arise. In the novel environment, the genetic basis of flowing time is indeed different (Fig 1/2), supporting that adaptive evolution acts on an alternative inheritance system. However, we found that plasticity and plastic alleles were highly correlated with environment factors (Figure 5 b/c). This highlights that plasticity is not an independent trait of the phenotype mean and should be constrained by selection on phenotype mean. Due to the mean effects, plastic alleles may already be under selection before environment changes occurs. Overall, our results shown that plasticity could be adaptive and it does not challenge the classical theory of nature selection and provide a broader view of the evolutionary process.

## Materials and methods

### Data

Two public datasets with measured mean flowering time from four to eight replicates using

514 *Arabidopsis thaliana* worldwide accessions (Alonso-Blanco *et al*., 2016; Exposito-Alonso *et al*., 2019) were reanalysed in this study. Flowering time scored as days to flowering were downloaded from the supplementary tables (Alonso-Blanco *et al*., 2016; Exposito-Alonso *et al*., 2019). Imputed SNP matrix was obtained from http://1001genomes.org/data/GMIMPI/releases/v3.1/SNP_matrix_imputed_hdf5/1001_SNP_MATRIX.tar.gz. After filtering for minor allele frequency greater than 0.03 and pairwise r2 smaller than 0.99, 1,396,438 SNPs were kept for downstream analysis. 19 climate variables were downloaded from the world climate database (http://www.worldclimate.org)

### Estimating kinship heritability and genetic correlation

*Kinship* heritability was estimated by fitting the following mixed linear model.

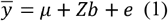

Here, 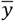 is flowering time mean of 514 lines. *μ* is the intercept representing the population mean. *Z* is the corresponding design matrix that satisfies *ZZ*^*T*^ = *G*, where *G* is the genomic kinship matrix estimated from GCTA (Yang *et al*., 2011). Therefore, *b* follows 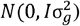. *e* is the residual with 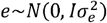. Kinship heritability is calculated as interclass correlation 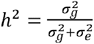. All the analysis were performed in GCTA (Yang *et al*., 2011). Pairwise genetic correlation among FT measured under the ten environments was estimated by a multivariant mixed linear model in ASReml.

### Quantification of plasticity measurements

We quantified flowering time plasticity using two types of metrics. The first type is called specific measurements (SP), describing the flowering time plasticity of two specific environments. SP measurements were calculated as all pairwise difference in flowering time value from individual environments, resulted in 45 measurements in total. The second type is overall plasticity (OP), describing flowering time plasticity across all environments. The OP measurements were calculated in three ways. First, across environment variance in rank transformed phenotype from ten environments (VarR) (Vanous *et al*., 2019) was used. Second, flowering time mean measurements from ten environments were used to perform a principal component analysis and the first and second principal component were used (Yano *et al*., 2019). Third, we used Finlay–Wilkinson regression (FWR) (Finlay & Wilkinson, 1963; Lian & De Los Campos, 2016) to partition the flowering time mean into two components, one component being constant across all environments and a second component respond dynamically to environment changes.

### Genome-wide association analysis

Genome-wide association analysis was performed using the a linear mixed model implemented in GCTA (Yang *et al*., 2011). SNPs with p values smaller than 0.05/Me (Me is the number of independent SNPs estimated using methods described in Li et al(Li *et al*., 2012)). Effect sizes of detected QTL were extracted from GCTA output. Conditional analysis, implemented in the “cojo” module of GCTA, was performed to screen for additional association signals at each QTL region. The effect sizes b of each SNP were estimated by the above mixed linear model.

### Partitioning flowering time mean and plasticity to contribution from QTL and polygenic effects

We used the following mixed linear model (Model 1) to partition the flowering time mean to contribution from QTL and polygenic effects.

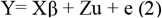

Y is a vector of 514 trait mean/plasticity of each individual (genotype) at each environment. *e* is the normally distributed residual. and u is a random effect vector of the polygenic scores for the 514 individuals. Z is the corresponding design matrix obtained from a Cholesky decomposition of the kinship matrix G, estimated using the genome-wide markers, excluding the detected QTL using GCTA (Yang et al. 2011). The Z matrix satisfies ZZ’=G, therefore, 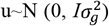. e is the residual variance with 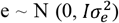. *Xb* is fixed effect, representing contribution from QTL. *X* is a design matrix, which contains QTL genotypes of each line, and β are the corresponding effect sizes. Contribution from QTL and polygenic effects was calculated with 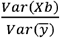 and 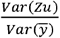.

### Testing interaction between QTL and environment

We fitted the following models to test interaction between QTL and environment:

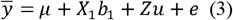

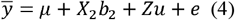

Since greenhouse condition is considerably different from that in common gardens, we only included the eight flowering time measurements from two common gardens in this analysis. 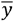 is a vector of length 514*8, representing concatenated FTm from 514 individuals and eight environments. *μ* is population mean, and e is normally distributed residuals. Both *μ* and e is of the same length as 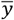. *u* is a random effect vector representing polygenic effects for the 514 individuals. The Z matrix is the corresponding design matrix. Z matrix satisfies *ZZ*^*T*^ = *G*⊗*I*, where I is an 8*8 identity matrix, therefore, u is normally distributed. *X*_1_ is a design matrix include genotypes from one QTL, common garden coded as factor, plant density coded as factor, and water availability coded as factor b_1_ is a vector containing the corresponding effect sizes. *X*_2_ is a design matrix similar with *X*_1_ but with one additional column as genotype times environment factor. Likelihood-ratio test were used to compare goodness of fit for these two models with and without the additional interaction term. This analysis is implemented in customised R script which is available upon request.

## Author Contributions

Y.Z. designed and supervised this study. Y.J., Y.H. collected the data and implemented the analysis. H.S. contributed to the statistical analysis. X.F, Q.C, W.Z., Z.L, Z.C R.H.and F.H. offered valuable input during the design and implementation of this study. Y.Z., Y.H. and H.S. wrote the manuscript.

All authors have read and approved the manuscript and have no conflicts of interest to declare.

## Acknowledgements

We thank Harry Wu and Örjan Carlborg for their constructive comments to improve the manuscript. We thank the 1001 Genome consortium for generating such a valuable resource, and specific thanks to Weigel Lab for releasing the flowering time datasets.

## Data availability

All phenotype measurements and metadata of samples are included in supplementary tables.

