## Supplemental Figures for "Polygenic basis of strong and rapid flowering time response to environment perturbations in wild *Arabidopsis thaliana* population"

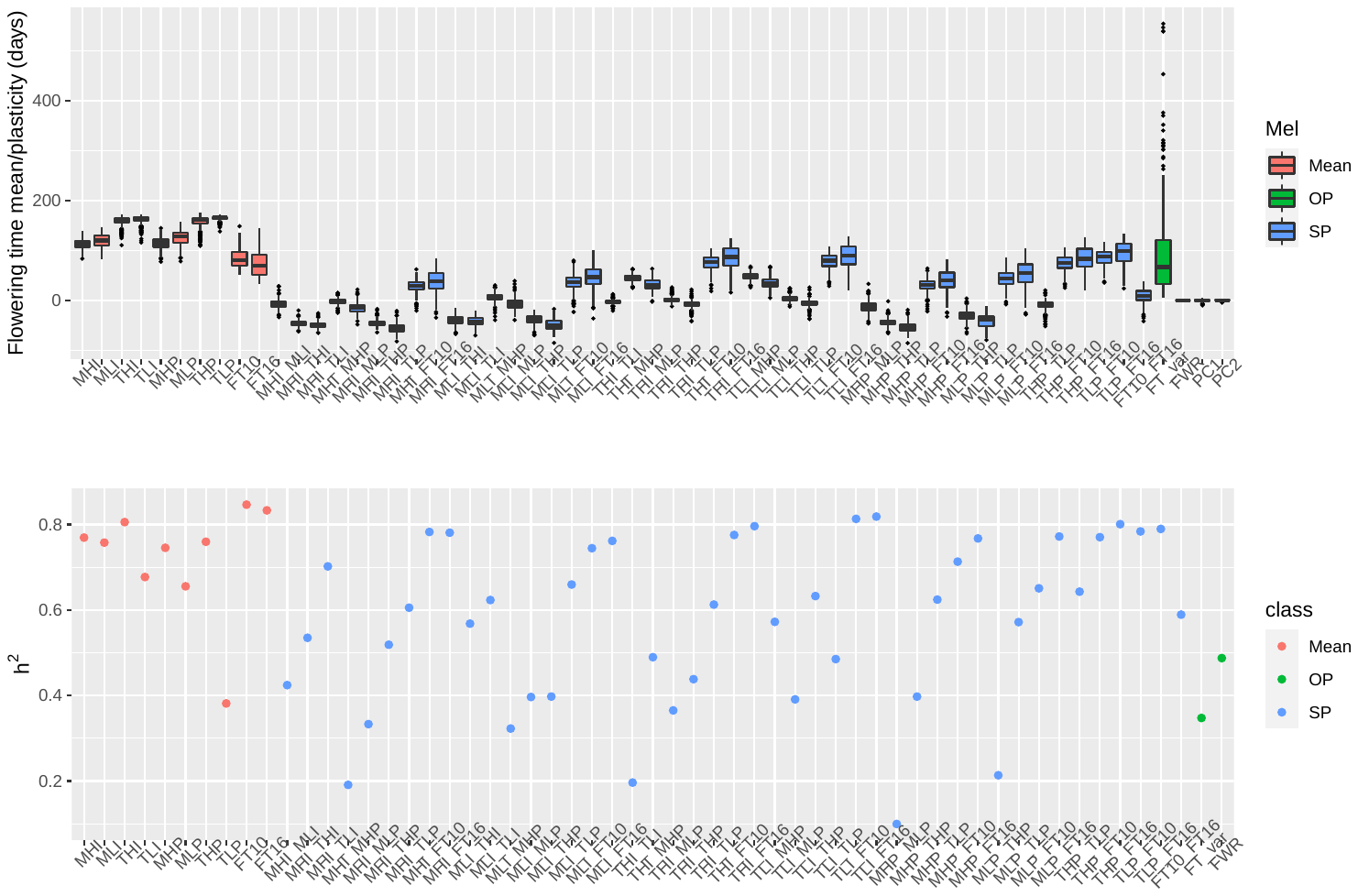

**Fig S1. Phenotype distribution and narrow sense heritabilities for the FTm and FTp measurements.** A) Boxplot of FTm and FTp measured at ten studied environments. B) Estimated narrow-sense heritability (h2) for FTm and FTp.

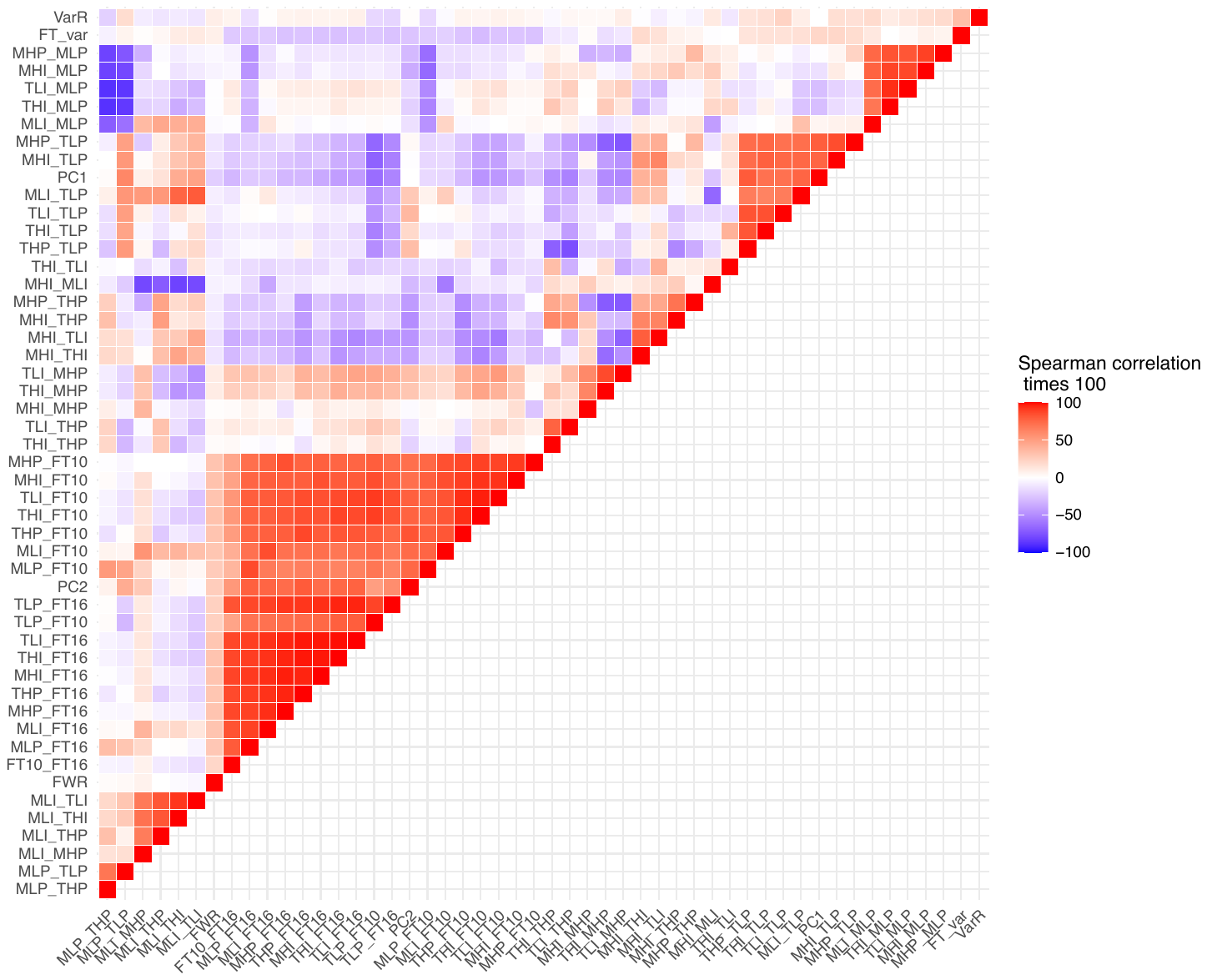

**Fig S2. Pairwise Spearman rank correlation among 48 plasticity measurements.**

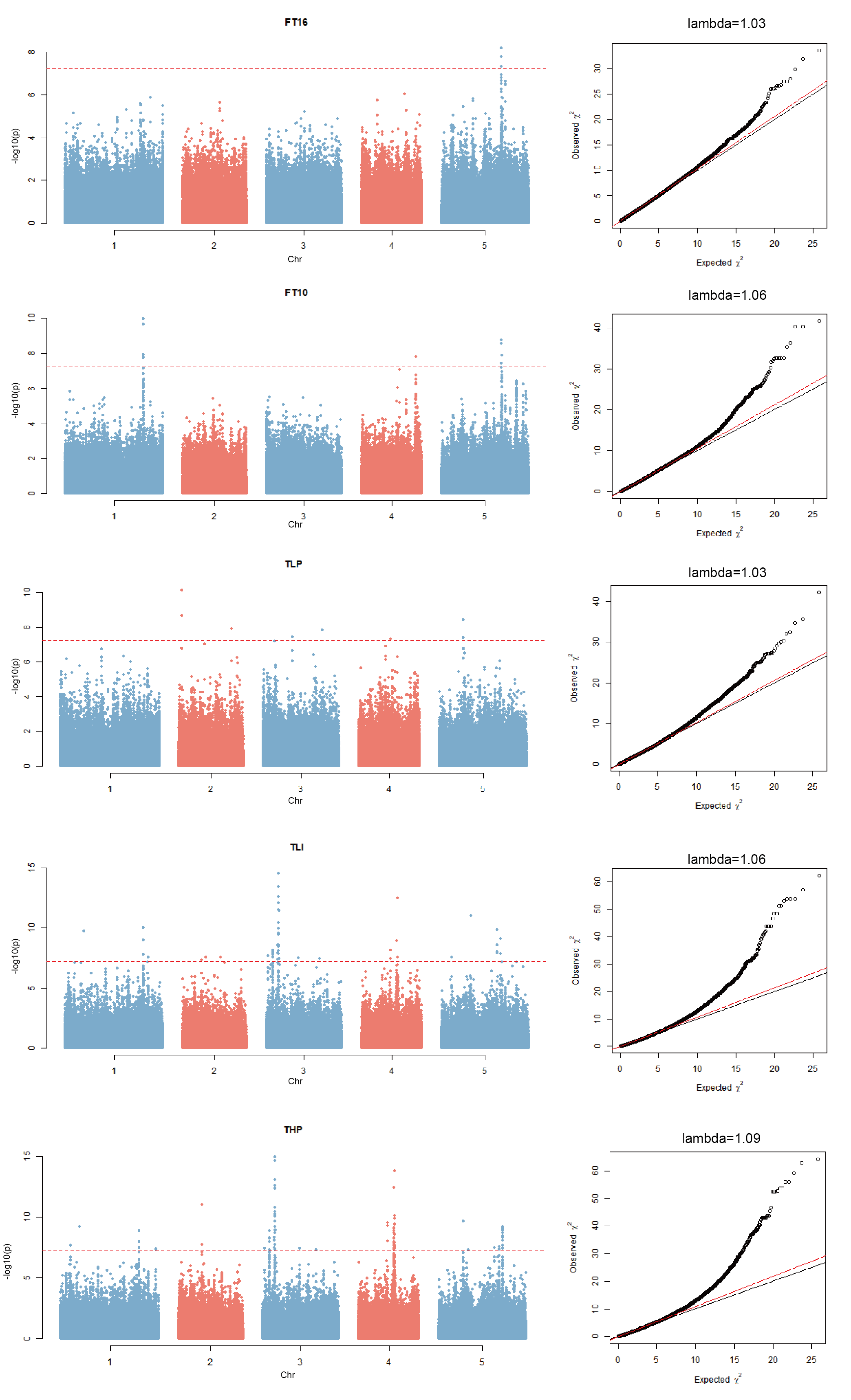

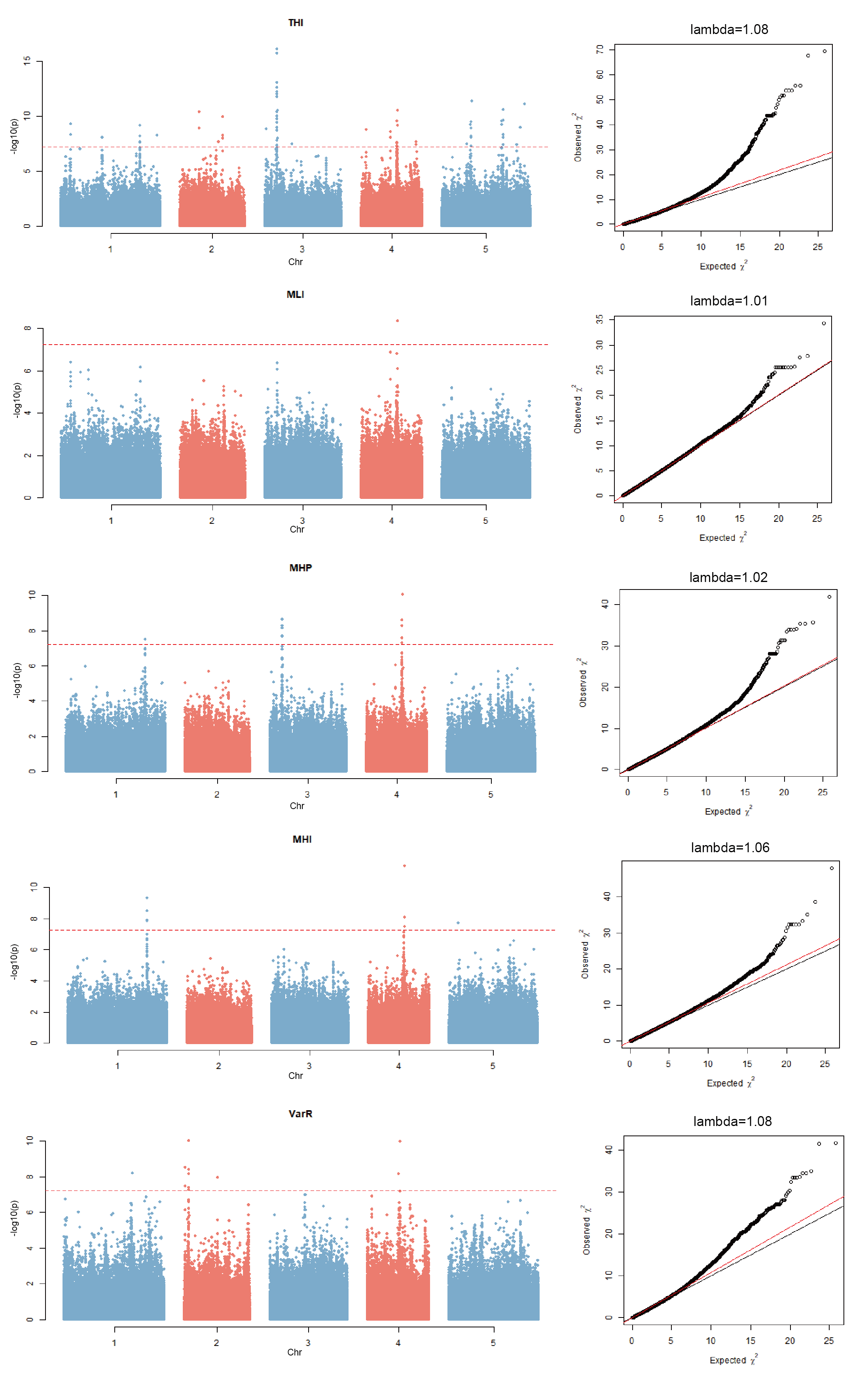

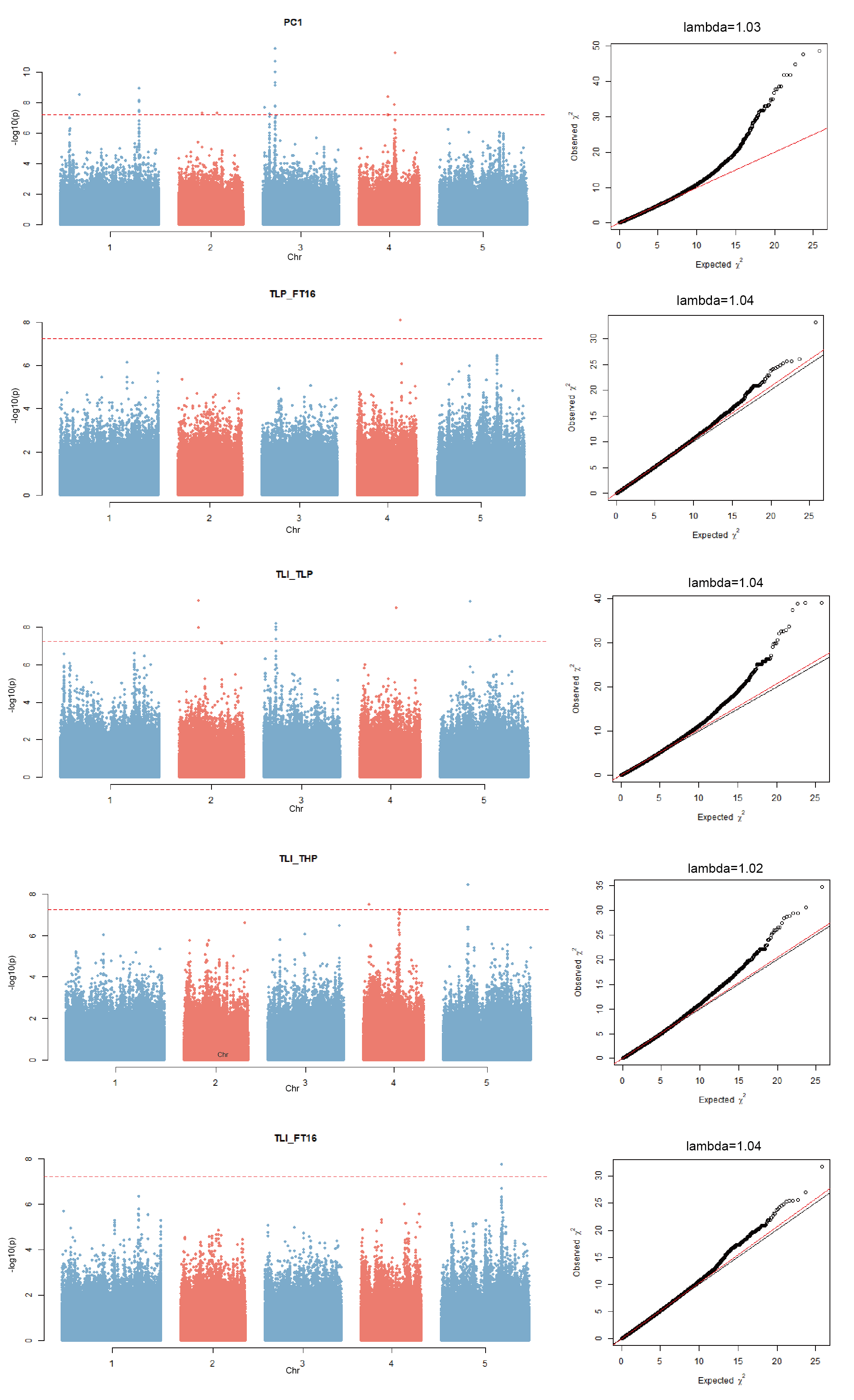

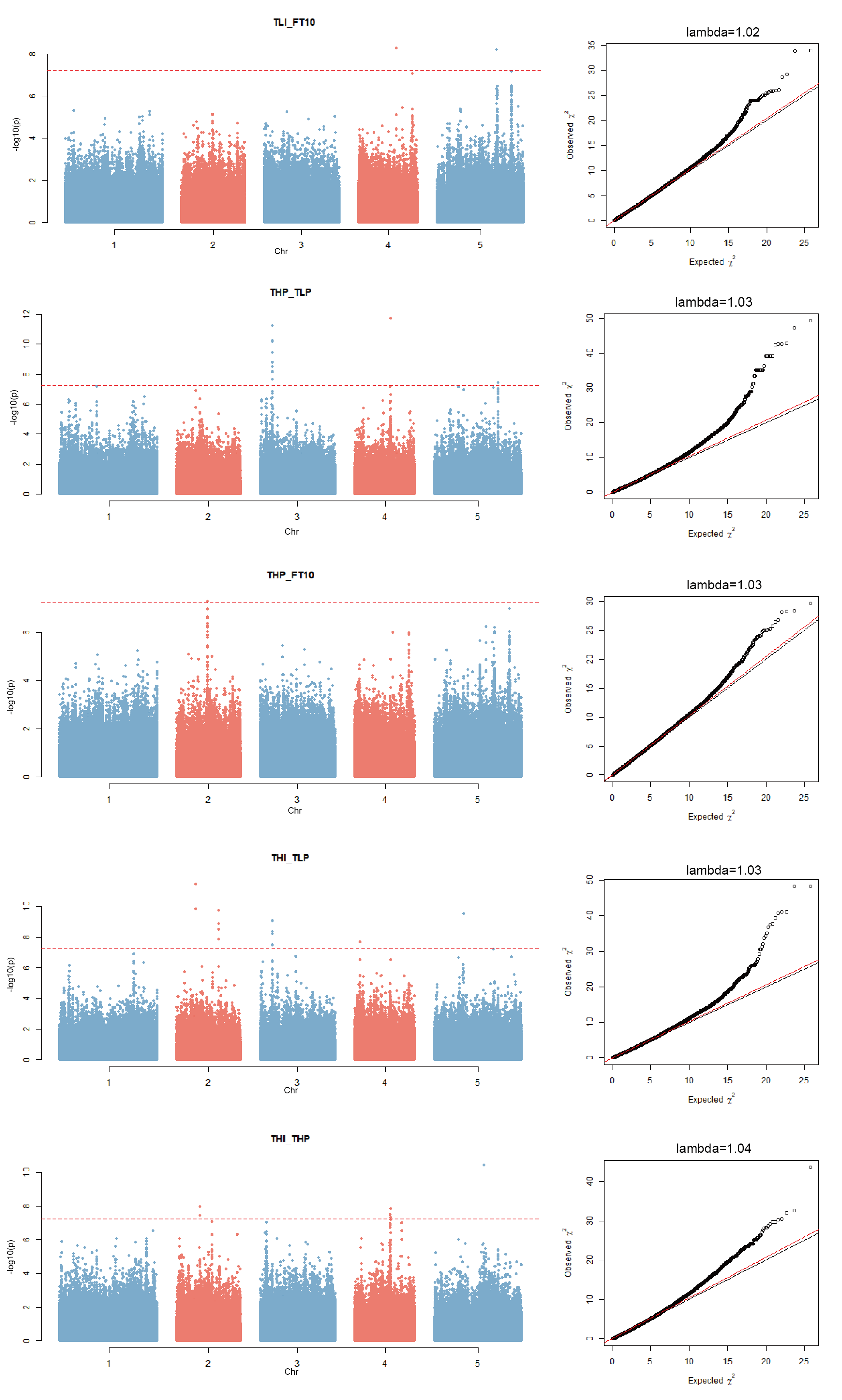

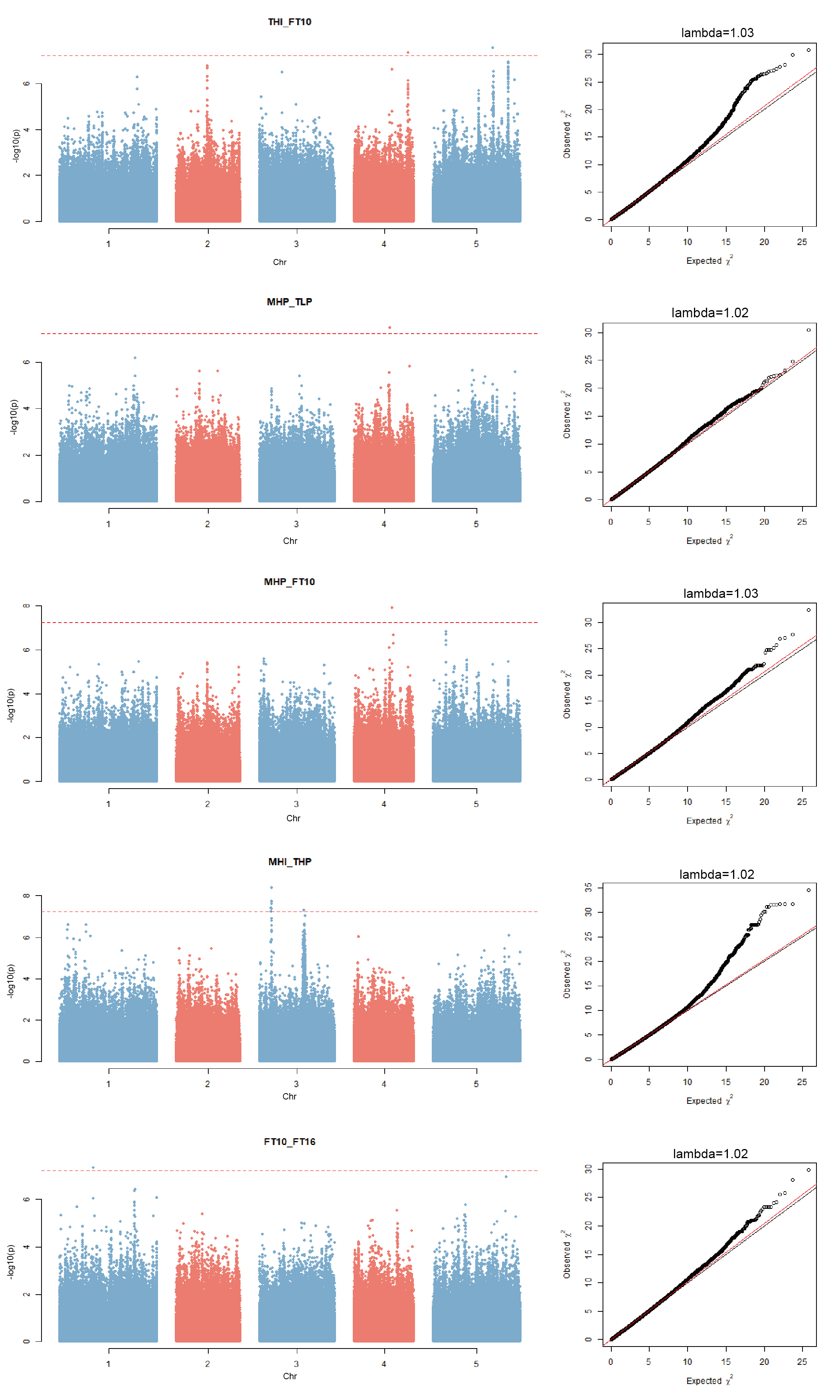

**Fig S3. Manhattan plots for FTm and FTp measurements for which significant association was detected.**
